# Single fiber proteomics of respiratory chain defects in mitochondrial disorders

**DOI:** 10.1101/421750

**Authors:** Marta Murgia, Jing Tan, Philipp E. Geyer, Sophia Doll, Matthias Mann, Thomas Klopstock

**Affiliations:** Department of Proteomics and Signal Transduction, Max-Planck-Institute of Biochemistry Am Klopferspitz 18, 82152 Martinsried, Germany; Department of Biomedical Sciences, University of Padova, Via Ugo Bassi, 58/B, 35131, Padua, Italy; Friedrich-Baur-Institute, Dept. of Neurology, University of Munich, 80336 Munich, Germany; NNF Center for Protein Research, University of Copenhagen, Copenhagen, Denmark; German Center for Neurodegenerative Diseases (DZNE), Munich, Germany; Munich Cluster for Systems Neurology (SyNergy), Munich, Germany

**Keywords:** Chronic progressive external ophthalmoplegia (CPEO), laser microdissection, mass spectrometry, mitochondrial disorders, proteomics, single muscle fibers

## Abstract

Mitochondrial DNA mutations progressively compromise the respiratory chain of skeletal muscle, resulting in a mosaic of metabolically healthy and defective fibers. The single fiber investigation of this important diagnostic feature has been beyond the capability of large-scale technologies so far. We used laser capture microdissection (LCM) to excise thin sections of individual muscle fibers from frozen biopsies of patients suffering from chronic progressive external ophthalmoplegia. We then applied a highly sensitive mass spectrometry (MS)-based proteomics workflow to analyze healthy and defective muscle fibers within the same biopsy. We quantified more than 4000 proteins in each patient, covering 75% of all respiratory chain subunits, and compared their expression in metabolically healthy and defective muscle fibers. Our findings show that mitochondrial disease causes extensive proteomic rearrangements, affecting the OPA1-dependent cristae remodeling pathway and mitochondrial translation. We provide fiber type-specific information showing that increased expression of fatty acid oxidation enzymes occurs in defective slow but not fast muscle fibers. Our findings shed light on compensatory mechanisms in muscle fibers that struggle with energy shortage and metabolic stress.

## Introduction

Mitochondrial disorders are multisystem diseases characterized by defects in the assembly and function of the mitochondrial respiratory chain. Mutations of both mitochondrial DNA (mtDNA) and nuclear DNA (nDNA) have been identified as leading causes of these disorders, summing up to a total prevalence of adult mitochondrial disease of 1 in 4,300 [1]. The human mitochondrial genome, which is maternally inherited, consists of 37 genes encoding 13 key proteins of the respiratory chain, 2 ribosomal RNAs and 22 mitochondrial transfer RNAs (tRNAs)[2]. The vast majority of mitochondrial proteins are encoded by the nDNA and the diseases caused by their mutations may be inherited in an autosomal dominant, autosomal recessive or X-linked manner [3]. Furthermore, age-associated neurodegeneration as in Parkinson’s and Alzheimer’s disease as well as aging itself has been associated with mitochondrial dysfunction [4]. Next generation sequencing has extended the repertoire and greatly improved the diagnosis of mitochondrial disorders [5]. However, despite this increasing amount of genetic knowledge, the pathophysiological mechanisms that contribute to the clinical manifestations of mitochondrial disorders are still poorly understood and diagnosis remains a challenge.

In most mtDNA-associated diseases, such as chronic progressive external ophthalmoplegia (CPEO), mitochondrial encephalomyopathy with lactic acidosis and stroke-like episodes (MELAS) and myoclonus epilepsy with ragged red fibers (MERRF), skeletal muscle shows a pathological mosaicism of metabolically compensated and decompensated fibers. These patients have heteroplasmy of healthy and mutated mtDNA and eventually reach a cell division and time-dependent threshold of molecular defects that trigger cell pathology. The mosaic is apparent using histochemical stains, typically the combined cytochrome c oxidase/succinate dehydrogenase (COX/SDH) staining. This is a common diagnostic test, whereby decompensated fibers are negative for the activity of complex IV of the respiratory chain, cytochrome c oxidase (COX-) but retain the blue (SDH) stain which reflects the activity of complex II of the respiratory chain (succinate–ubiquinone oxidoreductase, Fig. **1a**). Complex II is the only respiratory complex entirely encoded by nuclear DNA, therefore SDH stain is unaffected by deleterious mutations of mtDNA and a reliable marker of mitochondrial abundance. Compensated fibers are stained orange as a result of a functioning electron transport to complex IV (COX+, Fig. **1a**).

**Figure 1.**
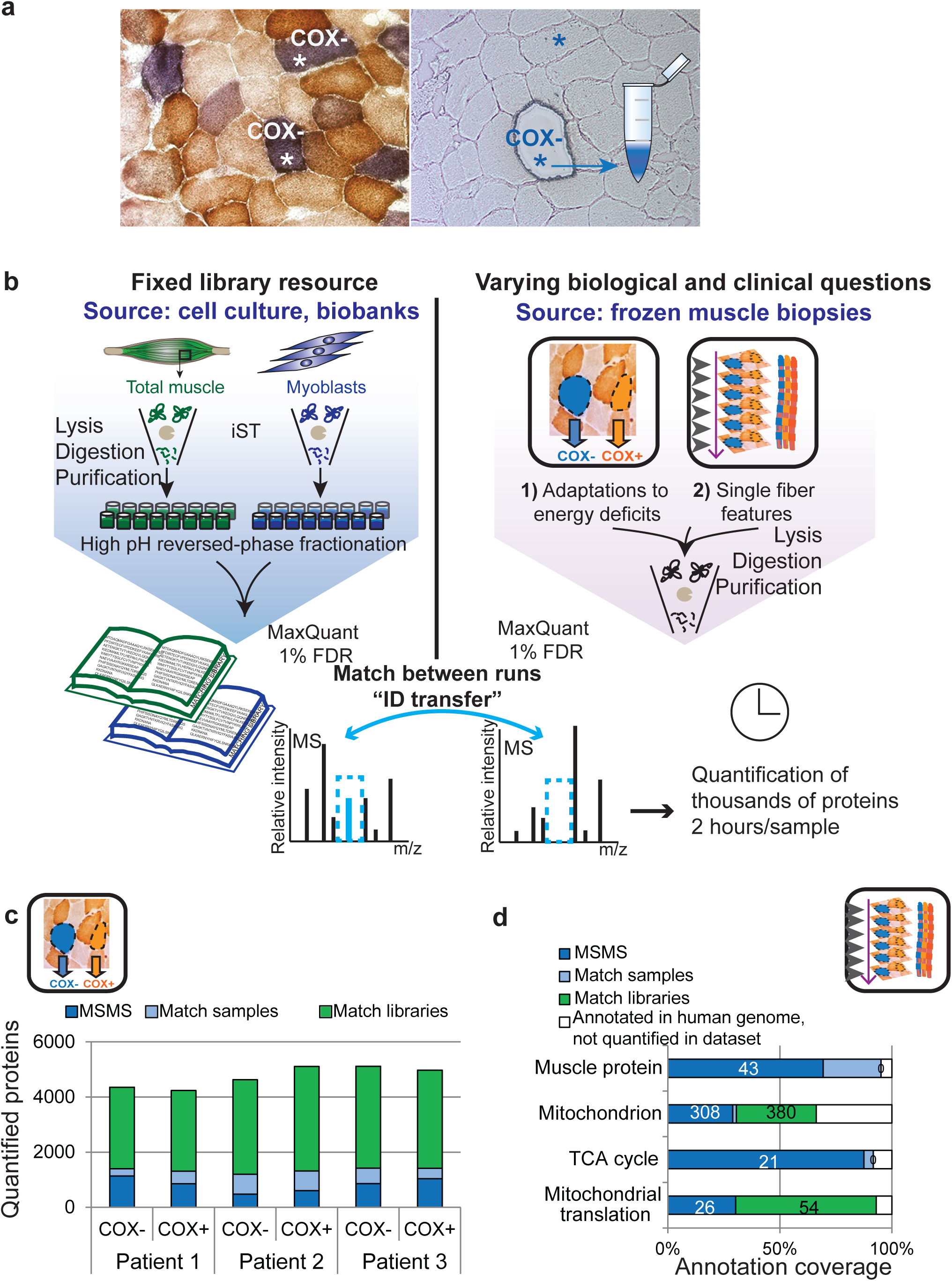
Design and workflow of the proteomic analysis of muscle biopsies from patients with mitochondrial diseases. (**a**) Schematic representation of laser capture microdissection applied here to COX/SDH staining. A stained section showing COX- and COX+ fibers is shown, the unstained serial section is used for fiber isolation. (**b**) Schematic representation of the proteomic workflow. Patient samples and lysates for the libraries are processed with a single vessel-based protocol, whereby all steps are performed in the same buffer. The peptides of the library samples are further fractionated prior to analysis. Peptides are measured by LC-MSMS, with a total duration of 2 hours/sample. Data analysis is performed by MaxQuant using the match between runs feature. (**c**) Number of proteins quantified in the dataset, illustrating the gain obtained by matching to the two peptide libraries (match libraries) or between samples only (match samples). (**d**) Coverage of different mitochondrial annotations (GO) in the single fibers dataset, expressed as percent of the corresponding terms annotated in the human genome (corresponding protein numbers are shown in the bar segments).

This pathological mosaicism is superimposed onto the physiological mosaic of different fiber types which characterizes skeletal muscle, one slow-type 1 and two fast-type 2 (2A and 2X), each with specific contractile and metabolic properties. In humans, slow fibers have more mitochondria than fast fibers.

In this study we asked how mitochondrial disease affects muscle fibers and how different fiber types cope with decreasing respiratory chain activity. The mosaicism of COX+ and COX-fibers offers the unique opportunity to investigate mitochondrial disease progression at the cellular level. To this aim, we generated a proteomic workflow for a rapid, robust and deep analysis of muscle fiber heterogeneity from patient biopsies, using laser capture microdissection (LCM) to separate COX+ and COX-fibers (**Fig. 1a**). Examples of a combination of LCM with proteomics exist in the literature in other types of myopathies, such as filaminopathies and desminopathies, and the proteome of intramyoplasmic aggregates has also been measured [6-8]. Exploiting recent advances in sample preparation and proteomic technology applied to skeletal muscle [9], we here quantify over 2000 thousands proteins from 20 individual fiber cryosections and over 4000 per patient (see Fig. S1a). This is the first single muscle fiber proteomic study in the context of mitochondrial disorders.

We address the adaptive molecular responses to a bioenergetic deficit at the level of individual fibers, uncovering fiber type-specific adaptive responses to disease, which could not be distinguished in a total lysate.

## Results

### Mass spectrometry-based proteomics workflow for muscle biopsies in mitochondrial disorders

We performed all sample preparations using the in-StageTip method [9] (see also Methods), which allows sample processing in a single reaction vessel to minimize sample loss, contamination, handling time and to increase quantification accuracy. The samples were measured in a quadrupole - Orbitrap mass spectrometer (see Methods) to obtain high sensitivity, sequencing speed, and mass accuracy[10]. We first built a fixed resource consisting of deep human skeletal muscle proteomes that we used as libraries of identified peptide features. To this end, we fractionated human muscle lysate and cultured myoblasts using a recently described loss-less nano-fractionator [9]. The peptide libraries allowed us to overcome the dynamic range problem driven by highly abundant sarcomeric elements which masks the identification of low abundant proteins. Using the “match between runs” feature of the MaxQuant analysis software[11, 12], we transferred identifications from the peptide libraries to the patients’ samples (**Fig. 1b**) [13]. The only pre-requisites for this strategy to be applicable to any muscle specimen and clinical need are identical liquid chromatography (LC) conditions and same biological species for all samples.

We show the scalability of the method from biopsy segments (as in the libraries) to single muscle fibers. With our workflow, triplicate proteome analyses of muscle sections from one patient can be carried out in 6 hours of total machine time and yield the quantification of over 4000 proteins (**Fig. 1c and S1a**). The same strategy allowed the proteomic analysis of serial sections of single fibers, in which we quantified 2440 +/-350 proteins on average (**Fig S1a and Table S1**).

We found that both libraries presented effective peptide separation into fractions with approximately 75% of all peptides being present in ≤ 4 fractions and up to 25% present in a single fraction (**Fig. S1b**). We quantified 5200 proteins (42000 peptides) in the muscle lysate, spanning over 7 orders of magnitude. The most intense quartile was specifically enriched in myosin and ATP synthase complexes and the least intense in exosome and RNA-processing proteins (**Fig. S1b and Table S2**). The myoblast lysate yielded 8600 proteins (74600 peptides), with gluconeogenesis and proteasome annotations enriched in the first quartile and DNA recombinase annotations in the last quartile ranked by intensity. The higher proteome coverage in myoblasts compared to muscle lysate is likely an effect of the lower dynamic range in their proteome (**Fig. S1c-d, Table S3,** see also [13]).

Our library-based strategy boosted protein identification with respect to the direct sequencing of the TopN method by tandem mass spectrometry (MSMS). The advantage of this method strategy was proportionally stronger in the dataset obtained with LCM of single muscle fibers, where peptides from low abundant proteins presumably fall below the intensity limit for MSMS identification (**Fig. 1c, d**). The gain in identification provided using libraries was sufficient to obtain 65% coverage of mitochondrial annotations. Abundant mitochondrial proteins, such as those of the respiratory chain and TCA cycle, were mostly identified by MSMS. However, the identification of potentially disease-relevant protein classes, such as those annotated as mitochondrial translation, reached high coverage (93%) only through matching to the libraries (**Fig. 1d**).

To assess quantitative reproducibility, we first analyzed several samples in technical triplicates. This resulted in excellent Pearson correlation coefficients of 0.98 (**Fig. S2**). The proteomes of pooled COX+ and COX-muscle fibers of the same individual (yellow squares) showed higher correlation (0.88-0.95) compared to those of different individuals (0.76-0.86). Correlations were on average lower in single fibers.

### Expression of respiratory complexes in COX+ and COX-muscle fibers

We set out to depict the mosaic of metabolically decompensated COX- and compensated COX+ fibers identified by COX/SDH staining in muscle biopsies of patients with CPEO. Using laser microdissection to cut individual fibers from 10 μm sections, we obtained separate pools of 100 COX+ and 100 COX-fibers from each of three different patients (**Fig. 1a, Table S4).** Using pools of muscle fiber sections helps avoid sampling biases, such as different fiber type composition leading to different mitochondrial content (**Table S5**).

In our dataset, we quantified 73 out of 95 proteins annotated to the respiratory chain complexes and ATP synthase in humans (GO annotation). We subdivided the subunits of the respiratory chain complexes based on their gene localization in mtDNA- and nDNA, respectively, then analyzed their differences in COX- and COX+ fiber pools. All COX complex IV subunits were more abundant in COX+ as compared to COX-fibers (**Fig. 2a**), serving as a proof of concept that the proteomic approach reflects the diagnostic histochemical discrimination. This expression difference was particularly evident when analyzing the subunits of COX-complex IV which are encoded by mtDNA. We observed this for all respiratory chain subunits of mtDNA origin (**Fig 2**, red boxes). Accordingly, COX+ muscle fibers of CPEO patients have been shown to contain more copies of mtDNA than the COX-counterparts[14]. Complex I subunits were also significantly higher in COX+ than in COX-fibers (**Fig. 2b**), while complex III and ATP synthase subunits showed similar expression levels in COX+ and COX-fibers (**Fig. 2c, d**). Three out of four subunits of SDH-complex II, a histological marker of mitochondrial content (see Fig. 1a), were more abundant in COX-fibers (**Fig. 2e**). In line with this finding, citrate synthase, a classical marker of mitochondrial mass, was 36% higher in COX-fibers, (**Fig S3a**). Increased mitochondrial biogenesis has been shown in healthy carriers of another mitochondrial disease, Leber’s hereditary optic neuropathy (LHON), and could thus be a compensatory response to energy stress conditions [15]. We could quantify three proteins involved in mitochondrial biogenesis in both COX+ and COX-fibers. They resulted more abundant in COX-compared to COX+ fibers (**Fig. S3b**). The assembly factors of mitochondrial complexes, were slightly higher in COX-fibers as compared to COX+ fibers in all complexes (**Fig. S3c**). Our data thus suggest that COX-fibers, despite increasing mitochondrial biogenesis and respiratory complex assembly as a possible compensatory mechanism for the energy deficit, have overall lower expression of respiratory complexes than COX+ fibers.

**Figure 2.**
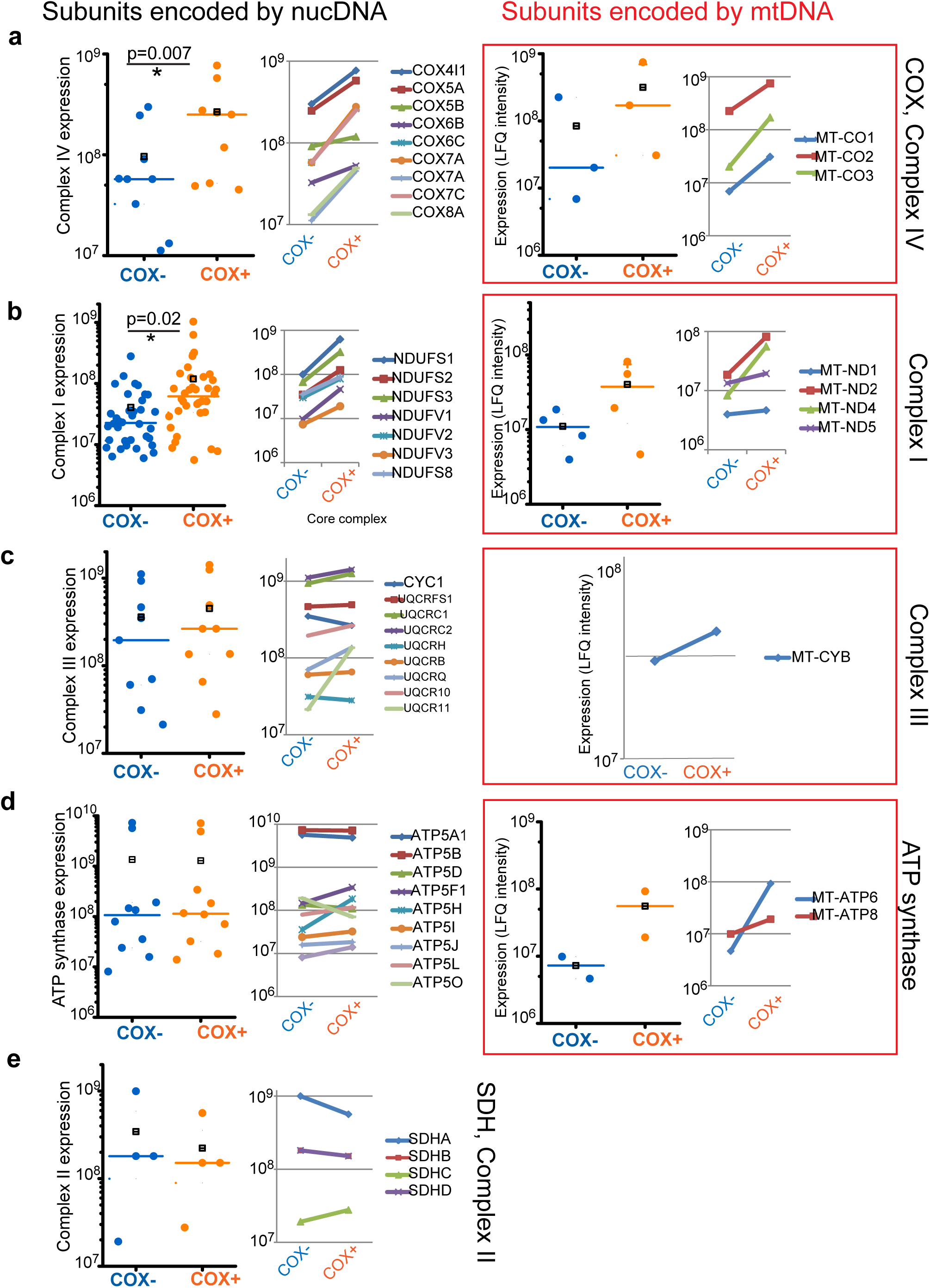
Expression of respiratory chain complexes in COX+ and COX-fibers. The subunits of each complex were subdivided into two groups according to their gene localization in nDNA (left column) or mtDNA (right column, red box) and the expression of all subunits was compared using scatter plots in COX-(blue) and COX+ (orange) fiber pools. Median values are indicated by a line and mean by a black line in a square. Significant differences are indicated by (*) and corresponding p value. Line graphs on the right of each scatter plot detail the expression of individual proteins of each complex in COX- and COX+ cells. (**a**) Expression of Complex IV (cytochrome oxidase) subunits. The median expression of each quantified protein of the complex in COX- and COX+ fibers is shown by the graph on the right. The values of the scale are Log_10_. (**b**) Expression of Complex I subunits. For simplicity, only the expression of the core catalytic complex is shown in the line graph. (**c**) Expression of Complex III subunits. (**d**) Expression of ATP synthase subunits. (**e**) Expression of complex II/SHD subunits, all encoded by nDNA. Each data point represents three patients in technical triplicates. All represented as in (a).

### Adaptive molecular responses to mitochondrial dysfunction at the cellular level

A comparison of the proteomes of COX+ and COX-fiber pools from all patients, revealed 580 proteins with significantly different expression (p<0.05). Principal component analysis (PCA) of the significant proteins showed a separation of the COX+ and COX-fiber pools along component 3 (**Fig. 3a**). This was driven by a specific enrichment of proteins annotated to the respiratory chain (p<10^−7^) in COX+ fibers. In contrast, COX-fiber pools displayed a high expression of the fatty acid binding protein FABP5 and of Wolframin (WFS1), an endoplasmic reticulum (ER)-transmembrane protein involved in calcium homeostasis (**Fig. 3b**).

**Figure 3.**
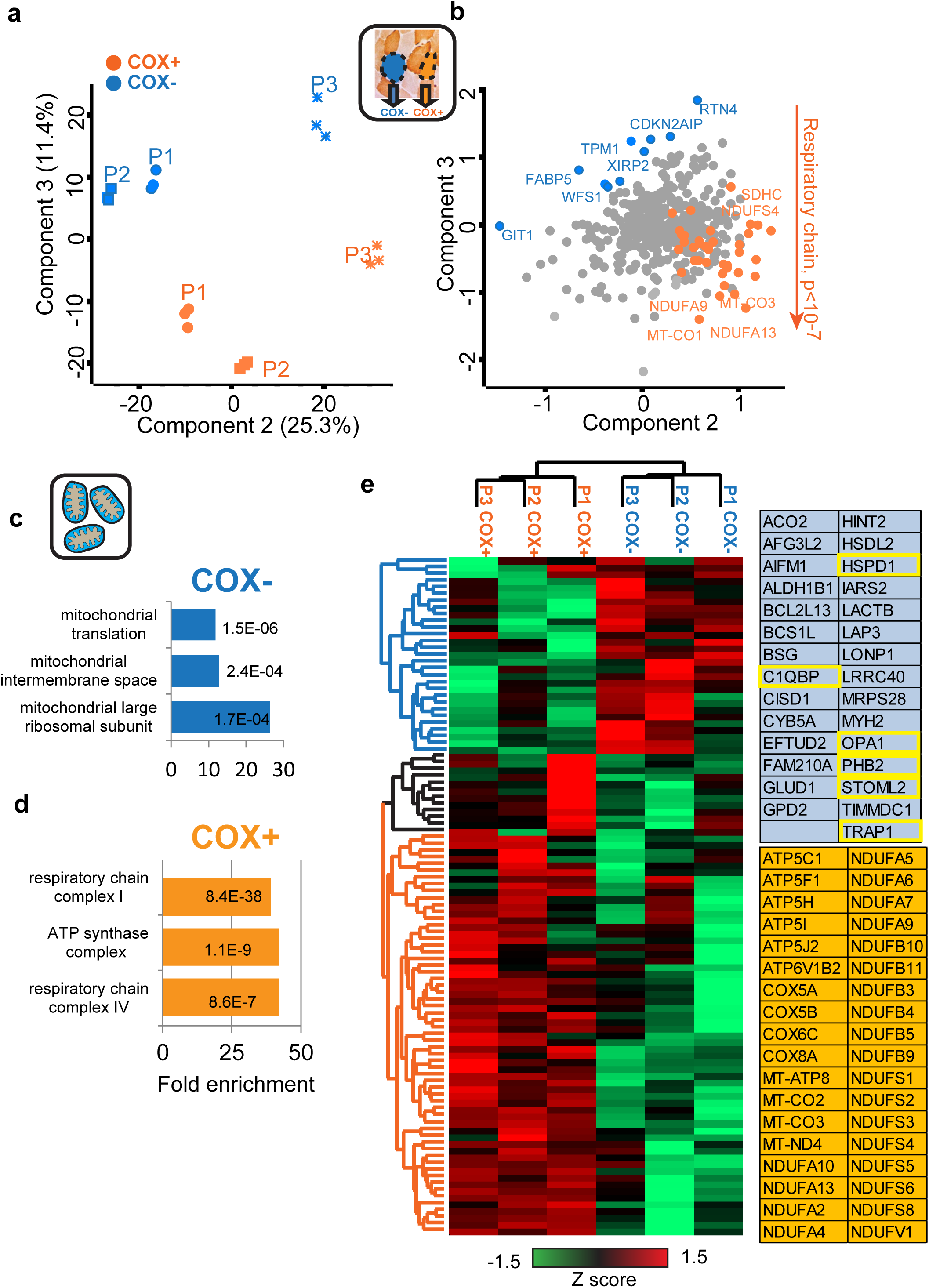
Proteomic features of COX+ and COX-muscle fiber pools. (**a**) Principal component analysis of mitochondrial proteins of three CPEO patients showing separation of COX+ (orange) and COX-(blue) fiber samples along component 3. Technical triplicates are shown for each data point. (b) Corresponding loadings of the PCA showing a highly significant enrichment of proteins annotated as respiratory chain on the COX+ side along component three. (c) Bar graphs showing the fold increase of the top three annotation enrichments of mitochondrial proteins with significantly higher expression in COX- and COX+ fiber pools. (Fischer’s exact test, FDR 0.02). (d) Unsupervised hierarchical clusters of mitochondrial proteins (normalized by CS) with significantly different expression in COX+ and COX-fiber pools. Each sample is the mean of technical triplicates. The proteins present in the two main clusters are shown in matching colors on the right. Proteins discussed in the text are highlighted in yellow.

We then focused on the mitochondrial proteome, using Mitocarta 2 [16] to select the mitochondrial proteins from the dataset (see Methods section). After normalizing mitochondrial protein expression by CS expression in each sample we could analyze the features of the mitochondrial proteomes of COX+ and COX-fiber pools correcting both for systematic differences of mitochondrial content between patients and for sampling differences in the amount of mitochondria. The comparison of the mitochondrial proteomes of all COX+ and COX-fiber pools showed 148 significantly expressed proteins (p<0.05). Next, we looked for annotations (GO and Keywords) that were significantly enriched in the protein subset with higher expression in COX+ fibers (106 proteins) and in COX-fibers (42 proteins) (**Table S6**). The former showed highly significant, >40-fold enrichments in numerous annotations comprising the respiratory chain and electron transport (**Fig. 3c**). Proteins with higher expression in COX-fiber pools displayed significant enrichments (> 25-fold) in annotations related to mitochondrial translation (**Fig. 3d**). This might be a general compensatory mechanism for the defective expression and function of the respiratory chain in COX-fibers (see Discussion).

Based on unsupervised hierarchical clustering of the mitochondrial proteins with significantly different expression between COX+ and COX-fibers, we revealed two main clusters consisting of COX+ and COX-fibers. The top cluster was specifically enriched in the annotation ‘phosphoprotein’ (p<10^−4^) and included proteins such as STOML2, PHB2 and OPA1 involved in cardiolipin binding and inner membrane biogenesis, cristae remodeling, mitochondrial fusion and respiratory supercomplex assembly, as well as the mitochondrial chaperones TRAP1 and HSPD1. The second cluster consisted of proteins that are up-regulated in the COX+ fibers and significantly enriched (p<10^−7^) in annotations related to oxidative phosphorylation and electron transport (**Fig. 3e**). Our data support the notion that COX-fibers upregulate key proteins controlling the mitochondrial network organization, possibly in an attempt to enhance their reduced bioenergetic efficiency.

### Comparison of the mitochondrial proteome of individual CPEO patients

To elucidate the effects of mitochondrial disease on the proteome of each patient we analyzed the mitochondrial proteome of the three CPEO patients individually and compared protein expression between COX+ and COX-fibers. In each patient, PCA showed a clear separation of the two fiber pools.(**Fig. 4a**). In all individuals, the separation was driven by a highly significant enrichment in components of the respiratory chain in COX+ fibers (p<10^−9^) while the drivers of the separation in COX-fibers were very heterogeneous (**Fig. 4b**). These proteins are of potential clinical relevance, as they may be part of the compensatory strategy used by the muscle fibers of a specific patient to respond to mitochondrial malfunctioning.

**Figure 4.**
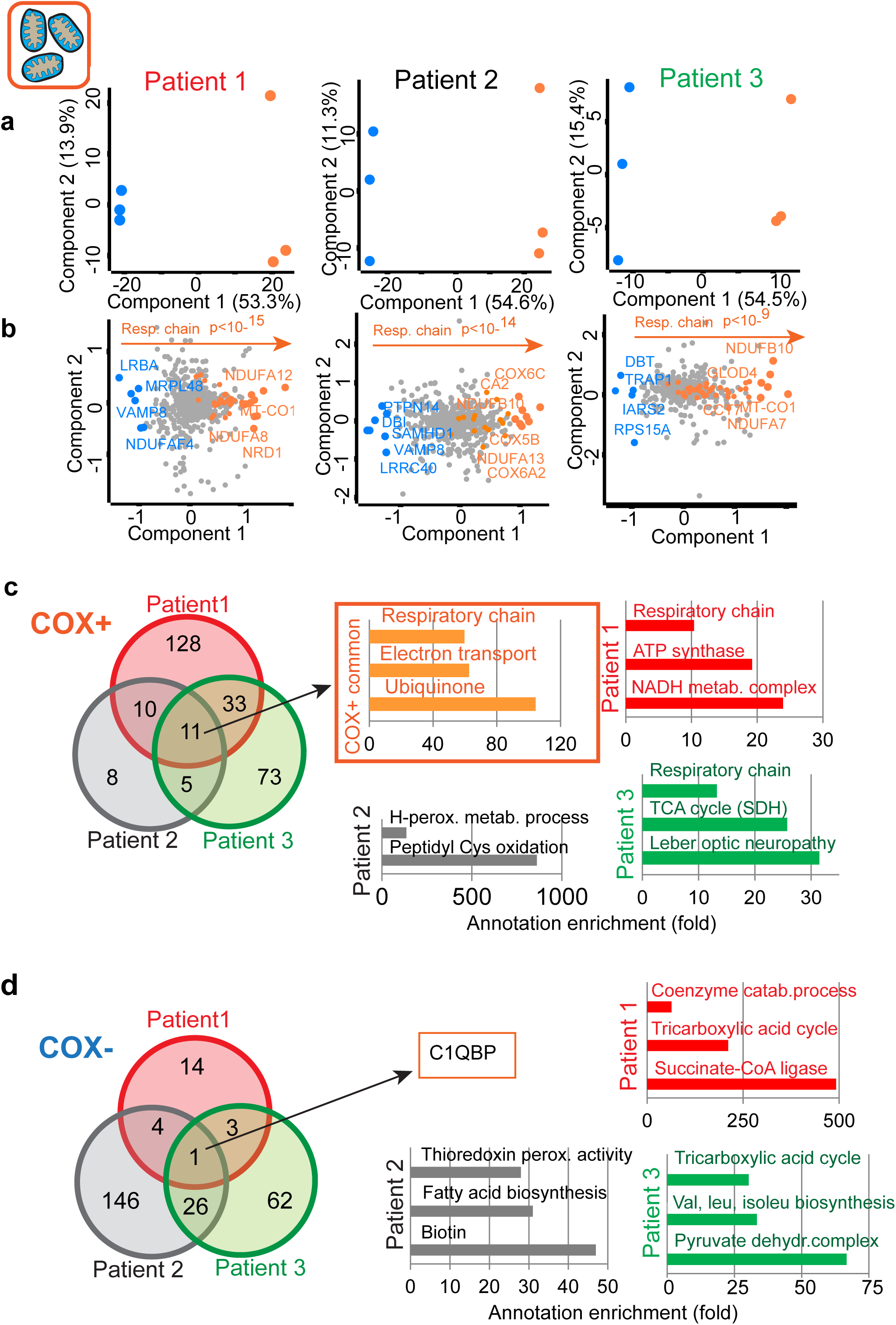
Patient-specific and shared features of mitochondrial disease. (**a**)Separation by principal component analysis of COX+ (orange) and COX-(blue) fiber pools of each patient (in triplicates). Separation occurs along component one, which defines the largest difference in the dataset. (b) Corresponding loadings driving the separation, with annotation enrichments and p values indicated by arrows. (c) Venn diagram showing common and patient-specific proteins that were significantly higher in expression in COX+ fibers. Highest annotation enrichments in both common and unique proteins (Fisher’s exact test, FDR 0.02), are shown with the bar graphs. (d) Analysis as in (c) for COX-fibers.

We performed for each patient a T-test comparing COX+ and COX-fiber pools to highlight the significant differences in their mitochondrial proteome. This analysis retrieved 182, 34 and 122 proteins with significantly higher expression in the COX+ fiber pools of patients 1, 2 and 3 respectively (**Table S7**). We then constructed a Venn diagram with these protein lists, highlighting 11 proteins that were common to all three patients and over 50-fold upregulated in annotations pertaining to the respiratory chain and electron transport (**Fig. 4c**). The same analysis was carried out on the proteins unique to each patient that were upregulated in COX+ fibers. This highlighted various annotations of energy metabolism in patient 1 and 3. Patient 2 showed overall fewer significant proteins in this analysis, with a unique enrichment in the annotation cysteine oxidation.

We then focused on the proteins with a significantly higher expression in COX-fibers, amounting to 22, 177 and 92 in the three patients (**Table S8**). Interestingly, only one protein, component 1 Q subcomponent-binding protein (C1QBP/p32), was commonly upregulated in COX-fibers of all three patients. C1QBP is a ubiquitously expressed protein localized predominantly in the mitochondrial matrix, whose function is poorly characterized. About 70% of proteins resulting from the t-test as significantly overexpressed in COX-fibers was specific for one patient only (**Fig. 4d**). The three annotations with largest enrichments were thus different among the patients. A common feature of patient 1 and 3 was the enrichment in the TCA cycle (**Fig. 4d**). A higher expression of these metabolic enzymes might be an indication of an increased usage of the Krebs cycle to alleviate the malfunctioning of the respiratory chain. TCA cycle upregulation is not evident in patient 2. Clinical, biochemical and genetic heterogeneity leading to such metabolic variations are likely causing different responses to therapeutic interventions.

### Impact of mitochondrial disease at the single fiber level

Human slow-type 1 fibers are characterized by an oxidative metabolism and more mitochondria than the fast-type 2A and 2X fibers, which are characterized by a higher expression of glycolytic enzymes [17]. Comparing pure fiber types can thus more precisely pinpoint the specific disease pathogenesis at the cellular level. To this aim, we cut 40 cross-sections of 10 µm each from muscle biopsies of the 3 CPEO patients, histochemically stained every second section as a morphological reference, and excised 20 serial sections of individual muscle fibers using laser microdissection (**Fig. 1a)**. With this procedure, each single fiber encompasses 400 µm of tissue. The serial sections of the same fiber were pooled and processed together for single-shot MS analysis. We isolated three COX+ and three COX-single fibers from each patient. MS-based proteomics allowed us to directly quantify different myosin isoforms and thus determine fiber type (**Table S5**)[18]. Out of 18 single fibers analyzed, 13 were pure slow-type 1 fibers, as defined by the expression of more than 80% of MYH7, three were pure fast fibers expressing a majority of MYH2A and three were mixed 1/2A type (**Fig. S4a**). To confirm the precision of our fiber type assignment we selected the proteins involved in muscle contractions (Keyword annotation “muscle protein”), which are typically expressed in a fiber type-specific manner. An unsupervised hierarchical clustering of all pure single fibers divides the fibers assigned as fast from those assigned as slow. Among the proteins characterizing the two main clusters are myosin and troponin isoforms, precisely segregating into fast and slow gene products (**Fig. S4b**).

We next focused on the proteome of pure slow-type 1 fibers, analyzing the expression of mitochondrial proteins that were expressed in at least 9 of 12 slow fibers (217 proteins). We could separate COX+ and COX-slow fibers along component 2, which was driven by the significantly different expression of respiratory chain components (**Fig. 5a-b**). Among the drivers of the separation we highlighted several proteins which were higher in COX-slow fibers and play key regulatory roles in the architecture of the inner mitochondrial membrane.

**Figure 5.**
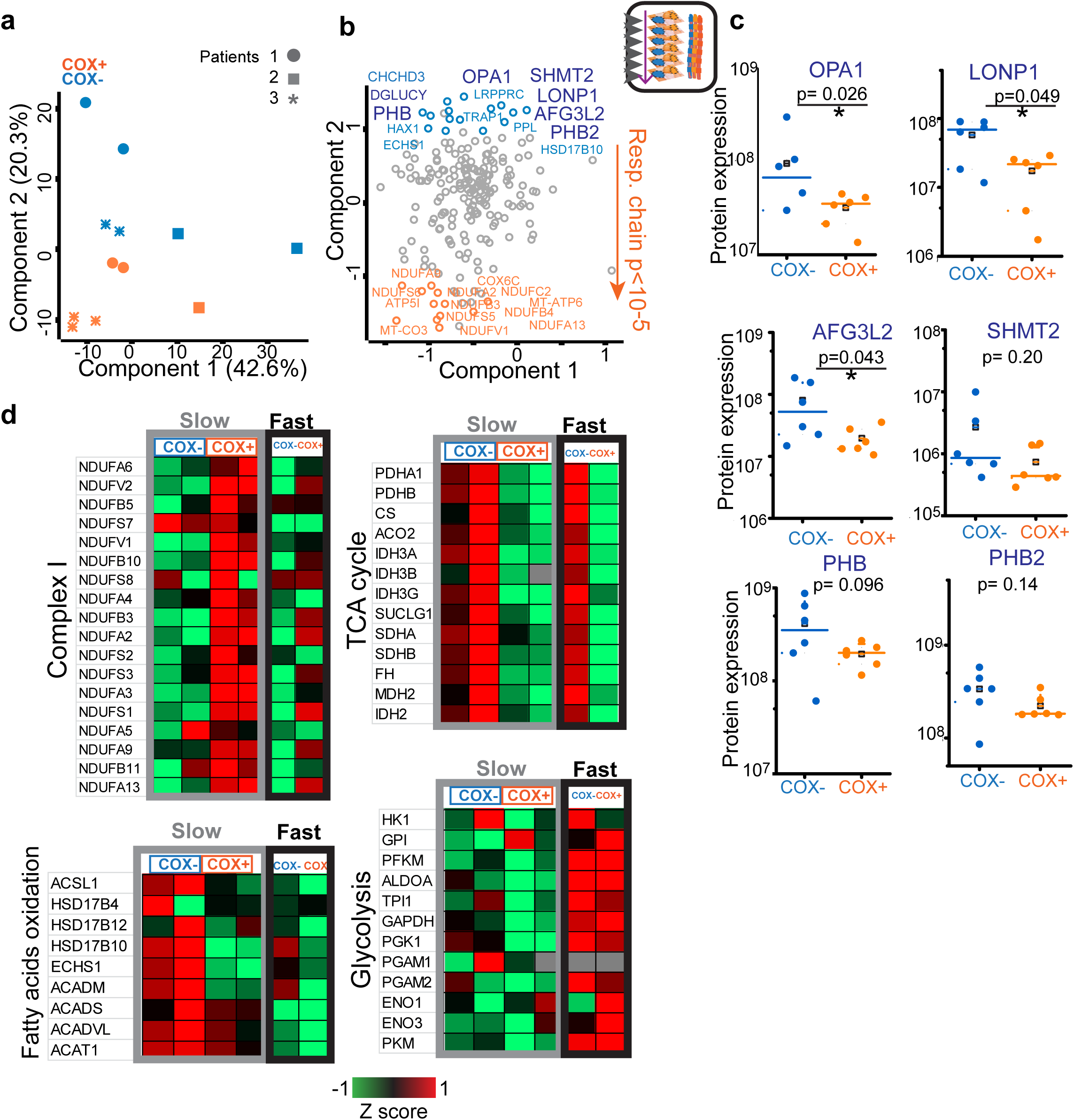
Single fiber features and fiber type specificity in mitochondrial myopathies. (a) Principal component analysis of the mitochondrial proteins of pure slow single fibers. The fibers from three patients segregate into COX+ (orange) and COX-(blue) along component two. (b) Corresponding loadings with significant enrichments indicated by the arrow. (c) The expression of proteins labeled in dark blue in the previous panel is shown as scatter plots for COX- and COX+ fibers. Median values are indicated by a line and mean by a black line in a square. Significant differences are indicated by (*) and corresponding p value. Significantly different samples are marked with a line and asterisk. All p values (COX+ vs COX-) are reported. (d) Heat maps showing normalized protein expression (Z score) in single fibers from one patient, four slow fibers in total (two COX+ and two COX- as indicated) and two fast fibers, one COX- and one COX+ respectively.

We compared the abundance of these proteins in single COX+ and COX-slow-type 1 fibers (**Figure 5c**) and found a significantly higher expression in COX- of the Dynamin-related GTPase OPA1, a key member of the mitochondrial contact site and cristae organizing system (MICOS) involved in cristae remodeling and fusion[19]. We also report a significant overexpression of AFG3L2, a mitochondrial AAA protease in COX-fibers which controls mitochondrial fragmentation and calcium dynamics [20], as well as of the Lon protease homolog LONP1, a master regulator of mitochondrial protein homeostasis. Two prohibitin family members, PHB1 and PHB2, scaffolding proteins of the inner mitochondrial membrane and regulators of respiratory supercomplex formation were also upregulated in COX-compared to COX+ slow fibers. COX-slow fibers also display a higher expression of the mitochondrial folate enzyme serine hydroxymethyltransferase 2 (SHMT2), controlling the expression of respiratory chain enzymes[21]. Interestingly, the protein DGLUCY/c14orf1559, a recently characterized a D-glutamate cyclase that converts D-glutamate to 5-oxo-D-proline is also significantly upregulated in COX-fibers (**Fig. 5c)** [22].

To compare the effects of mitochondrial disease in slow and fast fibers we focused on the four slow fibers and two fast fibers obtained from patient 1 (see Fig. S4). We selected different GO terms and analyzed the corresponding proteins that we could quantify in all four fibers using heat maps (**Fig. 5d**). As expected based on our results (see Fig. 2-4), both COX+ slow fibers had a higher expression of the respiratory chain complex I than the corresponding COX-fibers. This could be observed also in the two fast fibers, at generally lower expression levels. This difference in the expression of respiratory chain components is ultimately the net effect of the mitochondrial disease on mitochondrial function, devoid of individual variability and fiber type. We observed that COX-fibers, both slow and fast, upregulated the TCA cycle enzymes, indicating that the change in expression of this metabolic pathway occurs as a reaction to mitochondrial disease in a fiber type-independent manner. We then looked for potential fiber type-specific features and focused on beta-oxidation of fatty acids, as we have previously shown that this pathway has a higher expression in the mitochondria of slow fibers than in those of fast fibers [18]. Indeed, beta-oxidation enzymes showed markedly higher expression in slow that in fast single fibers. Strikingly, COX-slow fibers further upregulated the beta-oxidation pathway compared to COX+ fibers. This disease-dependent increase was comparably negligible in fast fibers.

## Discussion

Mitochondrial disorders, in a strict sense, are the consequence of mutations affecting the respiratory chain. They frequently affect skeletal muscle, while other tissues are involved in a disease-specific manner. The overall relationship between mitochondrial DNA mutations and clinical presentation is complex and poorly characterized at the molecular level. A diagnostic marker of many mitochondrial disorders is the appearance of ragged red fibers reflecting metabolic decompensation of muscle fibers above a certain threshold of mutant mtDNA (heteroplasmy). The resulting mosaic of differently compromised fibers is best visualized by COX histochemistry showing COX+ and COX-fibers. Deeper insight into the underlying molecular mechanisms is much needed to better understand the pathogenesis of mitochondrial disorders. To provide mechanistic details, this analysis needs a combination of fine tissue dissection and large-scale technologies applied to minute tissue amounts. Here, we combined LCM of thin 10 um sections of muscle fibers with a sensitive MS-based proteomic workflow featuring recent technological advances developed in our laboratory. Reaching the level of single muscle fibers is a key feature of our approach to mitochondrial disease, as skeletal muscles are composed of variable fractions of slow and fast fibers, which have different contractile properties, mitochondrial content and general metabolic features. After measuring the proteome of manually isolated single fibers from mouse limb muscles [18]and human muscle biopsies[17], we here show that LCM of frozen muscle biopsies can be used to measure the proteome of single fibers in the context of human disease.

Our proteomic approach combines single-shot measurements of patient samples with a fixed resource consisting of extensively fractionated peptide libraries. In this way, we increased peptide and protein identification and were able to interface them with a broad spectrum of molecular and diagnostic questions. Our workflow allowed the quantification of over 4000 proteins from <50ng of patient material in just 3 hours of measurement time, despite the dynamic range of the muscle fiber proteome driven by highly abundant sarcomeric proteins. This depth allowed us a detailed analysis of the muscle proteome, providing accurate quantification of all respiratory complexes and almost complete coverage of TCA cycle and mitochondrial translation. We could clearly detect a significantly higher expression of the components of the respiratory chain in COX+ than in COX-fibers, which is the net effect of mitochondrial disease on skeletal muscle. As a consequence and compensatory mechanism, we could measure an increased expression of proteins involved in mitochondrial biogenesis in COX-fibers. Reaching the fine tissue detail of COX+ and COX-fibers sections was instrumental in uncovering these changes, which would have been masked by heterogeneity at the whole tissue level.

Our results clearly show that COX+ and COX-fibers are significantly different at the proteome level. While expressing respiratory chain components at a significantly lower level than COX+, COX-fibers upregulate mitochondrial ribosomes and proteins involved in the control of translation (**Fig. 3c**). Among these proteins, the complement component 1 Q subcomponent-binding protein (C1QBP) was common to all patients analyzed (Fig. 4). This protein is required for functional mitoribosome formation and mitochondrial protein synthesis as shown in C1QBP-knockout (KO) mice[23]. Recently, mutations in C1QBP have been detected in patients with mitochondrial cardiomyopathy, multisystemic involvement and combined respiratory-chain enzyme deficiency[24].

At the same time, COX-fibers increase the expression of several mitochondrial chaperones and of stomatin-like protein 2 (STOML2), which organizes cardiolipin-enriched microdomains in the inner mitochondrial membrane and controls the assembly of functional respiratory supercomplexes[25]. While being in line with previous reports in cellular models of mitochondrial disease, our proteomic data now quantify the molecular changes induced by mtDNA mutation at the level of the direct targets of disease, the muscle fibers. Because the ultimate effects of mtDNA mutations will reflect on the metabolic profile of muscle fibers, it would be interesting to carry out a metabolomic analysis of COX+ and COX-fibers following the technological progress of this field.

A relevant feature of our proteomic approach to mitochondrial disease is the ability to analyze mitochondrial disease in individual muscle fibers, by following and cutting the same fiber across 20 serial muscle sections. Indeed, skeletal muscle has a e heterogeneous composition in slow-type 1 and fast-type 2 fibers, which have different mitochondrial content. In the pathological context of mitochondrial disorders, fiber type composition is superimposed onto the pathological process giving rise to the COX+ and COX-fiber mosaic. To reduce the variables causing this extreme heterogeneity we selected a pool of single muscle fibers that were type-1 slow, based on the expression of MYH7, the slow myosin heavy chain isoform (Fig. 5). With this approach we eliminated confounding effects of the heterogeneous muscle fiber type composition, revealing a coordinated increase of the OPA1-dependent cristae remodeling program in the mitochondria of COX-slow fibers. This pathway controls the tightening of the mitochondrial cristae, which results in higher respiratory efficiency and limits the production of reactive oxygen species and cytochrome c release[26]. This fiber type-specific analysis also revealed that mitochondrial folate enzyme serine hydroxyl-methyl transferase 2 (SHMT2) is specifically upregulated in COX-fibers. It has been recently shown that defects in this enzyme cause impaired expression of respiratory chain components by interfering with tRNA methylation and causing ribosome stalling[21]. Comparing four slow and two fast single fibers from the same patient, we showed that an upregulation of the TCA cycle enzymes occur in both fast and slow COX-fibers and likely represent a fiber type-independent compensatory mechanism for the energy deficit caused by mitochondrial disease. Only COX-slow fibers, however, upregulated the enzymes involved in fatty acids beta-oxidation. Our single fiber analysis has thus uncovered a fiber type-specific metabolic shift induced by mitochondrial disease.

It remains to be determined whether the combination of the observed compensatory mechanisms ultimately provides a relief from the energy imbalance caused by respiratory chain defect, or whether it contributes to the pathogenesis of the disease by causing proteotoxic stress and the mitochondrial unfolded protein response. Mechanistic studies of how defects in the assembly and function of the respiratory chain are communicated to the myonuclei will be needed to understand the complex progressive pathogenesis of mitochondrial disorders and provide a molecular basis for a long needed targeted intervention.

Altogether, our study shows that decreased expression of respiratory complex proteins in COX-fibers is accompanied by increased expression of mitochondrial translation components. In addition, we measure changes in many proteins of the OPA1-dependent cristae remodeling pathway, affecting respiratory supercomplex formation and bioenergetic efficiency. These extensive proteomic rearrangements involve crucial metabolic pathways and mitochondrial dynamics, opening new working hypotheses for the development of targeted therapies.

## Materials and Methods

### Patients

All of the muscle specimens and myoblast cells were obtained from the biobank of the German network for mitochondrial disorders (mitoNET) at the Friedrich-Baur-Institute. The research project was approved by the ethics committee of the LMU Munich (N. 198-15) and the study has been performed in accordance with the ethical standards laid down in the 1964 Declaration of Helsinki. Muscle biopsies were obtained for diagnostic purposes and written informed consent was obtained from patients or their guardians. For the study we randomly selected 3 patients with CPEO, defined by the pathognomonic clinical phenotype, the presence of ragged red fibers on muscle biopsy and detection of an mtDNA deletion (**Table S4**). Diagnostic fragments of the right deltoid or left quadriceps muscles were collected by open muscle biopsy. Then muscle specimens were frozen in liquid nitrogen immediately and stored at −80°C. The control myoblast cells used for the library were randomly selected from 3 patients with no known mitochondrial disorder.

### Histochemistry

### Tissue preparation for cryosectioning

Muscle specimens were transferred in liquid nitrogen to the cryostat. Alternate serial sections (10 µm) were adhered to Superfrost plus microscope slides for histochemical staining and to membrane slides for laser microdissection (Leica, No. 11505158). Superfrost plus slides were air-dried for about 24h and then stored at −20°C for the next histochemical staining. The membrane slides were stored at −80°C prior to cutting and processing for MS-based proteomics.

### Sequential cytochrome c oxidase / succinate dehydrogenase (COX/SDH) histochemistry

Slides were allowed to dry at room temperature for 10 min and processed according to standard protocol [27]. For COX staining, sections were incubated in cytochrome c oxidase medium (100 μM cytochrome c, 4 mM diaminobenzidine tetrahydrochloride, and 20 μg/ml catalase in 0.2 M phosphate buffer, pH 7.0) for 90 min at 37 °C. Sections were then washed in standard PBS, pH 7.4 (2 × 5 min) and incubated in succinate dehydrogenase (SDH) medium (130 mM sodium succinate, 200 μM phenazine methosulphate, 1 mM sodium azide, 1.5 mM nitroblue tetrazolium in 0.2 M phosphate buffer, pH 7.0) for 120 min at 37 °C. Sections were then washed in PBS, pH 7.4 (2 × 5 min), rinsed in distilled and dehydrated in an increasing ethanol series up to 100%, prior to incubation in xylene and mounting in Eukitt.

### Laser capture microdissection (LCM)

The procedure was carried out essentially as described by Koob et al[28]. The images of whole COX/SDH stained slides and sections of interest were acquired and stored using a Leica LMD 7000 System. Next, we observed the unstained serial sections under the microscope at various magnifications and compared with the pictures of stained sections individually. According to the recognizable histochemical features of COX+ and COX-cells, we determined the coordinates of their corresponding unstained cells and cut them by LCM. 100 COX+ and 100 COX-cells were dissected for each patient. Similarly, we selected 3 COX+ and 3 COX-single fibers with clearly recognizable histochemical features for each patient, then excised 20 COX+ or 20 COX-serial sections for each fiber separately. The whole procedure was precisely timed for each sample and carried out in less than 30 min at room temperature. Fiber sections were captured by cutting the region of interest onto the caps of 0.5ml Thermo-Tube, which were carefully closed and immediately frozen in liquid nitrogen at the end of the procedure. Samples were stored at −80°C until used. Protein amount was determined in serial dilutions of fiber section pools resuspended in 8 M urea containing 10 mM Tris-HCl, pH 8.5, measuring the fluorescence emission of tryptophan (excitation 280 nm, emission 350 nm).

### Sample preparation and high pH-reversed phase fractionation

Total muscle (60 mg) was crushed in liquid nitrogen using a mortar and pestle. Powdered muscle samples and myoblast cells (3x 10^6^ cells) were resupended in 310 µl (5 µl/mg) and 200 µl SDC reduction and alkylation buffer, respectively. Fiber sections were resuspended Samples were further boiled for 10 min to denature proteins[9]. The total muscle sample was further mixed (six times 30 seconds and cooled on ice in between) using a FastPrep^®^-24 Instrument (MP Biomedicals). Protein concentration was measured using the Tryptophan assay and 250 µg were digested overnight with Lys-C and trypsin in a 1:25 ratio (µg of enzyme to µg of protein) at 37 °C, under continuous stirring. On the following day, samples were sonicated using a Bioruptor (Diagenode, 15 cycles of 30 sec) and further digested for 3 h with Lys-C and trypsin (1:100 ratio). Peptides were acidified to a final concentration of 0.1% trifluoroacetic acid (TFA) for SDB-RPS binding and 40 µg of peptides were loaded on four 14-gauge Stage-Tip plugs. Peptides were washed first with wash buffers (P.O. 00001, PreOmics GmbH) using an in house made Stage-Tip centrifuge at 2000 x g. Peptides were eluted with 60 µl of elution buffer (80% acetonitrile / 1% ammonia) into auto sampler vials and dried using a SpeedVac centrifuge (Eppendorf, Concentrator plus). Peptides were resuspended in 2% acetonitrile / 0.1% TFA before peptide concentration estimation using Nanodrop. About 40 µg of peptides of each sample were further fractionated into 54 fractions by the Spider fractionator device (PreOmics) with a rotor valve shift of 90 s and concatenated into 16 fractions using high pH reversed-phase fractionation, as previously described[29].

### Liquid Chromatography Tandem Mass Spectrometry (LC-MS/MS) analysis

Nanoflow LC-MS/MS analysis of tryptic peptides was conducted on a Q Exactive HF Orbitrap (Thermo Fisher Scientific) coupled to an EASYnLC 1200 ultra-high-pressure system (Thermo Fisher Scientific) via a nano-electrospray ion source (Thermo Fisher Scientific). Peptides were loaded on a 50 cm HPLC-column (75 μm inner diameter; in-house packed using ReproSil-Pur C18-AQ 1.9 µm silica beads; Dr. Maisch). Peptides were separated using a linear gradient from 2% B to 20% B in 55 min and stepped up to 40% in 40 min followed by a 5 min wash at 98% B at 350 nl/min where solvent A was 0.1% formic acid in water and solvent B was 80% acetonitrile and 0.1% formic acid in water. The gradient was followed by a 5 min 98% B wash and the total duration of the run was 100 min. Column temperature was kept at 60°C by a Peltier element-containing, in-house developed oven.

The mass spectrometer was operated in “top-15” data-dependent mode, collecting MS spectra in the Orbitrap mass analyzer (60,000 resolution, 300-1,650 m/z range) with an automatic gain control (AGC) target of 3E6 and a maximum ion injection time of 25 ms. The most intense ions from the full scan were isolated with an isolation width of 1.5 m/z. Following higher-energy collisional dissociation (HCD), MS/MS spectra were collected in the Orbitrap (15,000 resolution) with an AGC target of 5E4 and a maximum ion injection time of 60 ms. Precursor dynamic exclusion was enabled with a duration of 30 s.

### Computational proteomics

The MaxQuant software (version 1.5.4.3) was used for the analysis of raw files. Peak lists were searched against the human UniProt FASTA reference proteomes version of 2016 as well as against a common contaminants database using the Andromeda search engine[11, 30]. Carbamidomethyl was included in the search as a fixed modification, oxidation (M) and phospho (STY) as variable modifications. The FDR was set to 1% for both peptides (minimum length of 7 amino acids) and proteins and was calculated by searching a reverse database. Peptide identification was performed with an initial allowed precursor mass deviation up to 7 ppm and an allowed fragment mass deviation 20 ppm. For the relative quantification of MYH isoforms, only peptides unique to each isoform were used for protein quantification in MaxQuant. The relative expression of each MYH isoform is calculated as percent of the summed intensity of the four adult isoforms (MYH1, MYH2, MYH4, MYH7). The mass spectrometry proteomics data have been deposited to the ProteomeXchange Consortium via the partner repository with the dataset identifier PXD010489.

### Bioinformatic and statistical analysis

The Perseus software (version 1.5.4.2), part of the MaxQuant environment[31], was used for data analysis and statistics. Categorical annotations were provided in the form of UniProt Keywords, KEGG and Gene Ontology. Mitocarta2 scores were provided as numerical annotations. Label free quantification (MaxLFQ) was used for protein quantification in all experiments, using Z score where indicated[32]. When using pools of COX+ and COX-sections from individual patients, 100 sections were pooled, the peptides were purified and analysed by MS in technical triplicates. This was performed in three different patients (i.e. biological triplicates, each in technical triplicates, for COX+ and COX-respectively). For the single fiber analysis, three single COX+ and three single COX-fibers were isolated from each patient (i.e biological triplicates in three patients for COX+ and COX-respectively). We used t-test for binary comparisons and a p value of 0.05 for truncation. Normalization for mitochondrial content was performed dividing expression values of each sample by the corresponding expression of citrate synthase (CS). PCA and cluster analysis was performed in the Perseus software using logarithmic expression values of LFQ. For hierarchical clustering, LFQ intensities were Z-scored and clustered using Euclidean distance for column and row clustering. Where indicated, missing values were imputed by using random numbers from a normal distribution to simulate the expression of low abundant proteins. We used a width parameter of 0.3 of the standard deviation of all values in the dataset with a down shift by 1.8 times this standard deviation. Pathway enrichment analysis was performed using Fisher exact test with a Benjamini-Hochberg FDR cutoff of 0.02.

## Acknowledgements

We acknowledge all members of the Department of Proteomics and Signal Transduction, Nagarjuna Nagaraj and Stefano Schiaffino for help and fruitful discussions and are grateful to G. Sowa, Christian Deiml and I. Paron for technical support. We also thank Dr. C. Laub for his help with the Laser capture microdissection. We acknowledge the PRIDE Team for the data deposition. This work was supported by the Max-Planck Society for the Advancement of Science; The Louis Jeantet Foundation; the German Bundesministerium für Bildung und Forschung (BMBF) through the E-Rare project GENOMIT (01GM1603); and by European Union’s Horizon 2020 research and innovation program under grant agreement 686547 (MSmed project).

## Author contributions

TK, MMa, and MMu conceived the study and designed the experiments. JT performed stainings and laser capture microdissection. MMu, SD and JT prepared samples for proteomic analysis. SD and PEG performed peptide fractionation. MMu, SD and PEG performed MS analysis. MMu and MMa analyzed proteomic data. MMu and TK wrote the manuscript. All authors participated in discussion and revision of the manuscript. All authors read and approved the final manuscript.

